# Serotonin modulates nucleus accumbens circuits to suppress aggression

**DOI:** 10.1101/2025.11.26.690732

**Authors:** Zihui Zhang, Gavin C. Touponse, Pia J. Alderman, Teema Yassine, Matthew B. Pomrenze, Troy W. Harris, Amei N. Shank, Robert C. Malenka, Neir Eshel

## Abstract

Serotonin (5-hydroxytryptamine; 5HT) has long been considered anti-aggressive, but the mechanisms by which 5HT regulates downstream circuits to control aggression remain unclear. Combining fiber photometry, optogenetics, and miniaturized microscope recordings in double-transgenic mice, we find that 5HT levels ramp up in the nucleus accumbens during aggression, inhibiting a subset of D1 medium spiny neurons to suppress attacks. Our results reveal a novel 5HT-mediated neuromodulatory mechanism for limiting aggressive behavior.

## Introduction

Low levels of brain serotonin (5HT) have been associated with heightened aggression across the animal kingdom^1,2^. Drugs boosting the 5HT system, however, produce variable effects on aggression^3^, forcing clinicians to prescribe agents that sedate patients rather than specifically reduce aggression^4,5^. The complex behavioral effects of brain-wide 5HT modulation may be due to heterogeneity in the 5HT system, with distinct 5HT neuron populations projecting to different targets to mediate diverse behaviors^6–8^. While recent studies have begun to map specific 5HT targets onto their associated behaviors^8–11^, little is known about the target regions that control aggression, or how 5HT modulates neural activity in these targets^12,13^.

The nucleus accumbens (NAc) has long been identified as a key node controlling motivated behaviors^14,15^. Although most research has focused on the role of dopamine (DA) in this region^16^, recent studies have implicated NAc 5HT release in promoting a range of prosocial behaviors, including behavioral antecedents of empathy^17–20^. In particular, the medial shell subregion of the NAc, the site with the most abundant aggression-activated neurons^21^, is a major target of 5HT axons projecting from the dorsal raphe (DR)^10^. We therefore hypothesized that 5HT release in the NAc medial shell may also influence aggression, and that it would do so via distinct mechanisms from those engaged by DA.

## Results

### 5HT and DA levels in the NAc increase during aggression with distinct temporal profiles

To measure real-time changes in 5HT and DA levels in the NAc during aggression, we used fiber photometry to record either GRAB-5HT^22^ or GRAB-DA^23^ signals in the NAc during a resident-intruder assay^24^, where a weak intruder mouse was introduced to the home cage of the test mouse (**Fig. 1a, b; Supplementary Fig. 1a, b**). We found that 5HT levels were low at aggressive approach, ramped up during the attack itself, and peaked around aggression offset (**Fig. 1c-e**). While the peak and area under the curve (AUC) positively correlated with aggression duration (**Supplementary Fig. 2**), the slope of the 5HT ramp showed a strong negative correlation (**Fig. 1f**): the faster the rise in 5HT, the shorter the aggressive bout. In contrast, DA levels rose during both the pre-attack approach period and at attack onset (**Supplementary Fig. 1c-e**), and only weakly correlated with aggression duration (**Supplementary Fig. 1f**). These data suggest that DA and 5HT may signal different phases of aggression, with 5HT selectively engaged in mediating the termination of attacks.

**Figure 1:**
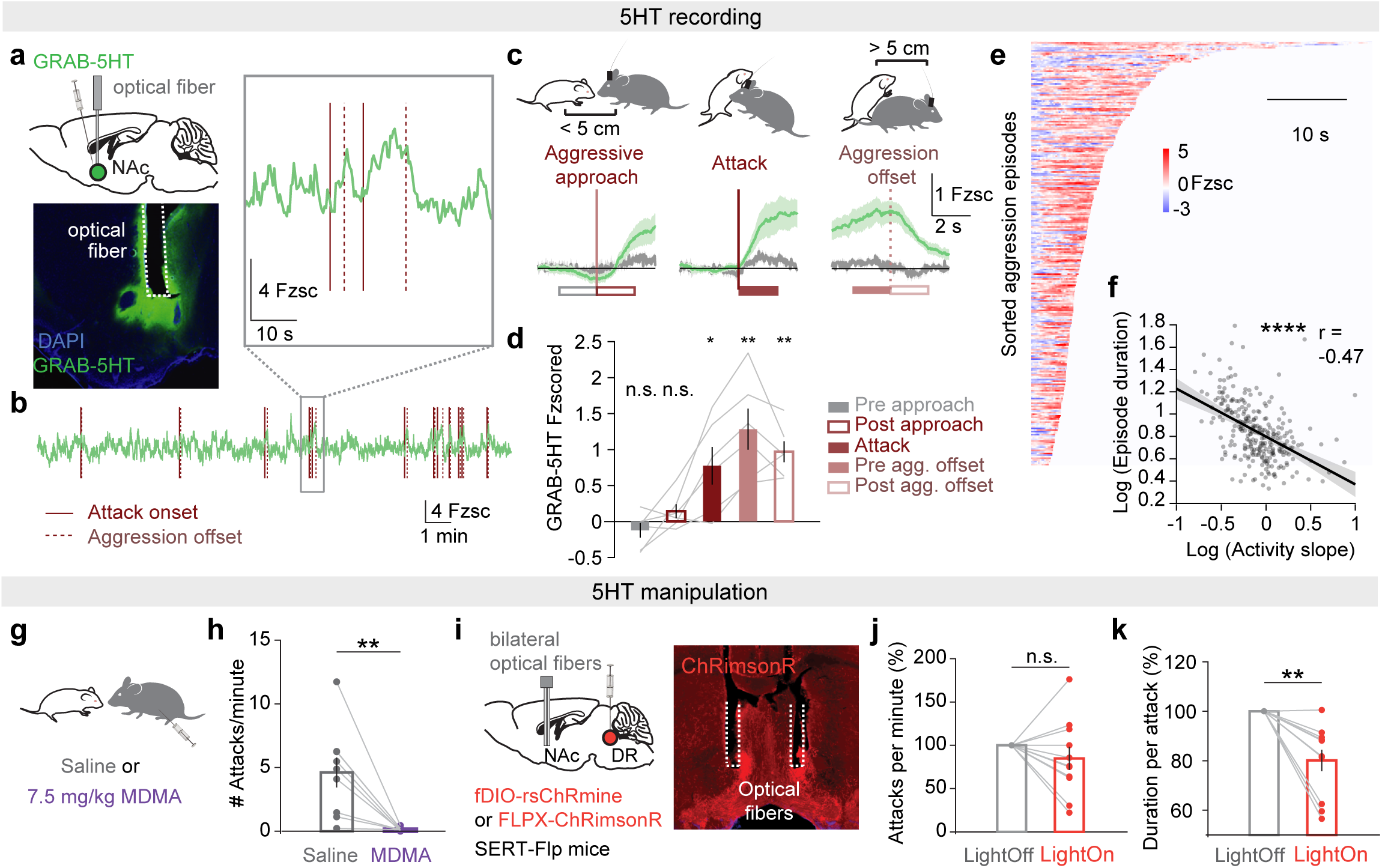
5HT signaling in the NAc during aggression. **(a)** GRAB-5HT virus was injected in the NAc for fiber photometry recording. **(b)** Example session of GRAB-5HT recording during a resident-intruder test. **(c)** Average GRAB-5HT (green) and control (grey) signal aligned to aggressive approach, attack, or aggression offset. **(d)** GRAB-5HT signal averaged in a 2-second window pre/post the behavioral events as marked in (c) (left to right: n.s. P = 0.28, 0.21, *P = 0.031, **P = 0.006, 0.001, one-sample t-test, n = 6 mice). **(e)** Heatmap showing GRAB-5HT signaling in individual aggressive bouts sorted by duration. **(f)** The duration of aggressive episodes negatively correlates with the slope of 5HT increase (***P<0.0001, r = −0.47, Pearson’s correlation coefficient). **(g)** Either saline or MDMA (7.5 mg/kg) was injected intraperitoneally 15 min before experiments. **(h)** Frequency of attacks in the resident-intruder test after injection of MDMA or saline (**P = 0.0039, Wilcoxon signed-rank test, n = 9 mice). **(i)** Flp-dependent excitatory opsin was injected into the DR of SERT-Flp mice. A bilateral optical fiber was implanted in the NAc for optogenetic stimulation of 5HT terminals. **(j)** Change in the frequency of attacks in photostimulation compared with non-photostimulation sessions (n.s. P = 0.19, Wilcoxon signed-rank test, n = 11 SERT-Flp mice). **(k)** Same as (j) but for the change in average duration of attack bouts (**P = 0.0020, Wilcoxon signed-rank test).

### 5HT but not DA release in the NAc suppresses aggression

We next tested the causal roles of 5HT or DA release in aggressive behavior. We initially examined the effects of (±)3,4-methylenedioxymethamphetamine (MDMA), a translationally-relevant compound that enhances both 5HT and DA release but promotes prosocial behaviors primarily by increasing 5HT levels in the NAc^18,25^. Consistent with prior studies^26^, administration of MDMA (7.5 mg/kg, i.p.) to the resident mouse 15 min before the resident-intruder test completely abolished aggression, compared to a saline control (**Fig. 1g, h**). Although this finding suggests that increased 5HT release may reduce aggression, the drug has numerous effects throughout the brain, and cannot be interpreted as selective to either 5HT or the NAc.

To more specifically manipulate either 5HT or DA release in the NAc, we injected the Flp-dependent excitatory opsin, FLPX-ChRimsonR or fDIO-rsChRmine, in either the DR of SERT-Flp mice (**Fig. 1i**), or the ventral tegmental area of DAT-Flp mice, and implanted optical fibers in the NAc (**Supplementary Fig. 1g**). Optogenetic activation of DA inputs in the NAc during the resident-intruder test did not affect either the frequency of attacks or the duration of each attack bout (**Supplementary Fig. 1h, i**), although we validated that photostimulation did trigger DA release (**Supplementary Fig. 1j-l**). In contrast, 5HT input activation significantly reduced the duration of attacks, without changing attack frequency (**Fig. 1j, k, Supplementary Fig. 3**) or basic locomotor, social, or valence-related behaviors (**Supplementary Fig. 4**).

To begin to determine the 5HT receptors mediating the anti-aggressive properties of NAc 5HT release, we infused a variety of 5HT receptor-specific agonists into the NAc immediately prior to the resident-intruder assay. Consistent with previous results examining the prosocial effects of NAc 5HT release^17^, infusion of a 5HT1b receptor agonist (CP93129) significantly reduced aggression, while 5HT1f and 5HT2c receptor agonists did not (**Supplementary Fig. 5**). These results suggest that in the medial NAc shell, 5HT release, but not DA release, is sufficient to reduce aggressive behavior, with a particular role in terminating attack bouts.

### D1 but not D2 MSNs in the NAc promote aggression

What downstream cells mediate the effects of NAc 5HT on aggression? The vast majority of NAc neurons are medium spiny neurons (MSNs) expressing one of two DA receptors: the D1 receptor, which is thought to augment neural activity in response to DA; or the D2 receptor, which does the opposite^27^. In the context of aggressive behavior, D1 MSNs in the NAc have been suggested to signal the rewarding aspect of aggression^21,28^. However, optogenetic stimulation of DA axons in the NAc, which presumably activates D1 MSNs, failed to affect aggression in prior studies^29,30^.

To determine how D1 and D2 MSNs are engaged during aggression, we injected the Cre-dependent calcium indicator, FLEX-GCaMP8m, in either D1-Cre or A2A-Cre mice, and recorded the activity of D1 or D2 MSNs during the resident-intruder test using a miniature microscope (**Fig. 2a, b**). On average, both MSN cell types exhibited increased activity during aggression (**Supplementary Fig. 6a-d**), but significantly more D1 MSNs than D2 MSNs showed robust activation across attacks (**Fig. 2c, d**). Examining individual trials, we found that the activity of D1 MSNs, but not D2 MSNs, positively correlated with the duration of aggressive episodes (**Fig. 2e, f**). This suggests that D1 MSN activity, like 5HT release, may influence the length of aggressive episodes. To test this hypothesis, we injected the Cre-dependent inhibitory opsin, NpHR3.0 in the NAc of either D1-Cre or A2A-Cre mice to optogenetically inhibit either D1 or D2 MSNs during the resident-intruder test (**Fig. 2g**). Similar to the effect of 5HT input activation, D1 MSN (but not D2 MSN) inhibition reduced the duration of attack bouts without influencing attack frequency (**Fig. 2h, I; Supplementary Fig. 3c-f**) or other locomotor, social, or valence-related behaviors (**Supplementary Fig. 6e-j**).

**Figure 2:**
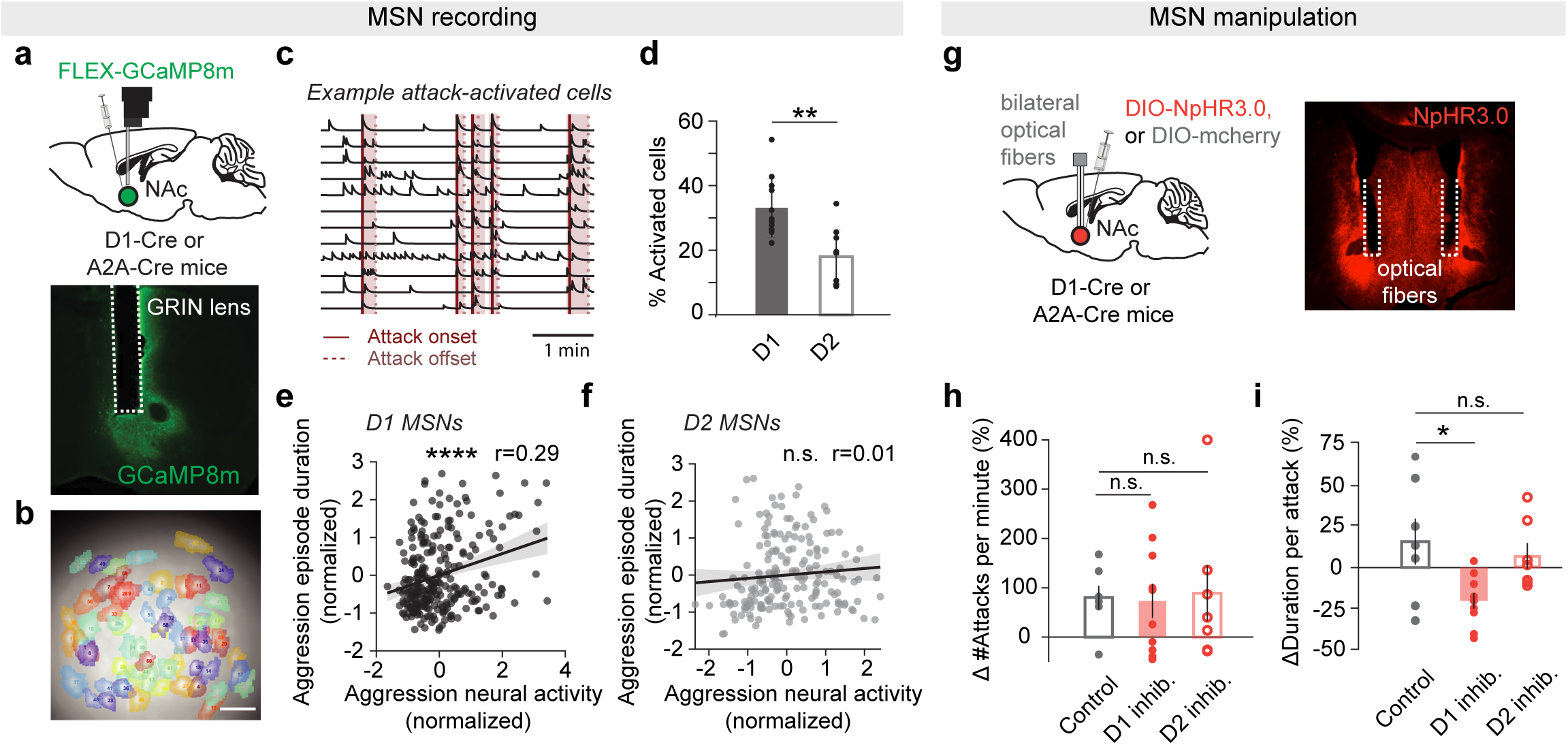
D1 and D2 MSN signaling in the NAc during aggression. **(a)** FLEX-GCaMP8m virus was injected in the NAc of D1-Cre or A2A-Cre mice for calcium imaging with a miniscope. **(b)** Example field-of-view showing detected neurons during the resident-intruder test (scale bar, 100μm). **(c)** Example calcium traces of cells robustly activated at attacks. **(d)** Percentage of D1 and D2 MSNs significantly activated during attacks (**P = 0.0011, Wilcoxon rank-sum test). **(e)** Correlation between the duration of the aggression episode and the average D1 MSN activity during aggression episodes (****P < 0.0001, r = 0.29, Pearson’s correlation coefficient). **(f)** Same as (e) but for D2 MSNs (n.s. P = 0.23). **(g)** Cre-dependent inhibitory opsin or control fluorophore (DIO-NpHR3.0 or DIO-mcherry) was injected in bilateral NAc of D1-Cre or A2A-Cre mice for optogenetic stimulation. **(h)** Change in the frequency of attacks in photostimulation compared with non-photostimulation sessions (n.s. P = 0.99, 0.96, one-way ANOVA followed by Tukey-Kramer test, n = 9 mice for D1 MSN inhibition, 7 mice for D2 MSN inhibition and 7 mice for controls). **(i)** Same as (h) but for the change in the average duration of attack bouts (n.s. P = 0.79, *P = 0.019, one-way ANOVA followed by Tukey-Kramer test).

### 5HT release in the NAc differentially modulates D1 and D2 MSNs

Given that 5HT stimulation and D1 MSN inhibition in the NAc produce similar anti-aggressive behavioral effects, we hypothesized that 5HT may specifically reduce the activity of D1 MSNs to suppress attack duration. To test this prediction, we first measured D1 or D2 MSN GCaMP8m activity using a miniscope when the mice were administered the same dose of MDMA (7.5 mg/kg) that abolished aggression in the resident-intruder tests (**Fig. 3a**). We found that MDMA administration led to a significant decrease in D1, but not D2 MSN firing (**Fig. 3b-d**). This result is consistent with our hypothesis that 5HT inhibits D1 MSN activity, but again does not control for other possible network effects of MDMA administration.

**Figure 3:**
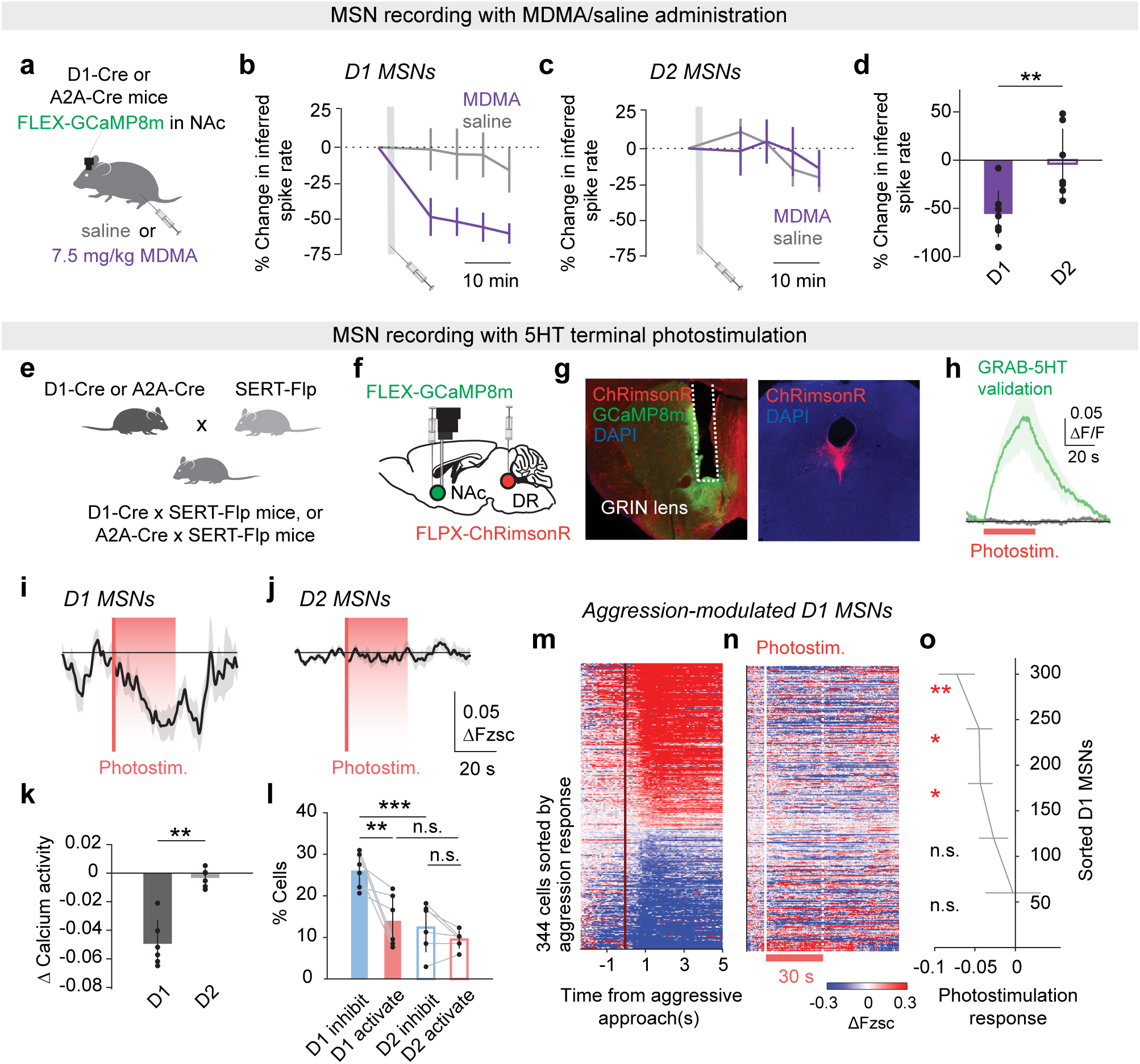
5HT preferentially inhibits D1 MSNs that signal aggression. **(a)** Experimental strategy: miniscope recording of either D1 or D2 MSNs expressing FLEX-GCaMP8m in the NAc after the mouse was injected with either saline or MDMA (7.5 mg/kg). **(b)** Change in the inferred spike rate in D1 MSNs after saline or MDMA injection. Grey areas indicate injection time. **(c)** Same as (b) but for D2 MSNs. **(d)** Change in the average inferred spike rate in D1 and D2 MSNs post MDMA injection compared to baseline (**P = 0.0037, Wilcoxon rank-sum test, n = 8 D1-Cre mice and 7 A2A-Cre mice). **(e)** D1-Cre or A2A-Cre mice were crossed with SERT-Flp mice to create double transgenic D1-Cre × SERT-Flp or A2A-Cre × SERT-Flp mice. **(f)** Double-transgenic mice in (e) were injected with Cre-dependent GCaMP8m virus in the NAc and Flp-dependent ChRimsonR in the DR. **(g)** Example brain slice showing the expression of GCaMP8m in NAc neurons and ChrimsonR in the terminals of 5HT neurons. **(h)** Validation experiment: SERT-Flp mice were injected with Flp-dependent ChRimsonR in the DR and GRAB-5HT in the NAc and an optical fiber was implanted in the NAc to record 5HT signals during photostimulation of 5HT terminals (n = 2 mice: 1 SERT-Flp × D1-Cre and 1 SERT-Flp × A2A-Cre). **(i)** Calcium activity of D1 MSNs aligned with photostimulation (30 s, 20 Hz, n = 6 D1-Cre, SERT-Flp mice). **(j)** Same as (i) but for D2 MSNs (n = 6 A2A-Cre, SERT-Flp mice). **(k)** Average calcium response to photostimulation in D1 and D2 MSNs (**P = 0.0022, Wilcoxon rank-sum test). **(l)** Percentage of cells robustly activated or inhibited across trials (n.s. P = 0.33, 0.86, **P = 0.0032, ***P = 0.00037, two-way ANOVA followed by Tukey-Kramer test). **(m)** Calcium activity of aggression-modulated D1 MSNs around aggressive approach. Each row is a cell, sorted by the calcium response in time 0-2 s (n = 344 cells, 12 sessions, 6 mice). **(n)** Calcium activity of the same cells in (m) aligned with photostimulation, which was delivered later that same day. **(o)** Photostimulation response of cells sorted by aggression response as in (m-n). (top to bottom: ** P = 0.007, *P = 0.035, 0.031, n.s. P = 0.35, 0.84, one-sample t-test).

To more specifically assess the effects of endogenous 5HT release on MSN activity, we generated double-transgenic mice by crossing D1-Cre or A2A-Cre mice with SERT-Flp mice, and injected them with Cre-dependent GCaMP8m virus in the NAc and Flp-dependent ChRimsonR virus in the DR (**Fig. 3e, f**). Placement of a GRIN lens over the NAc injection site allowed us to simultaneously stimulate 5HT release and record genetically-identified downstream neurons while mice rested in their home cages (**Fig. 3g**). Consistent with the MDMA results, optogenetic stimulation of 5HT inputs in the NAc (**Fig. 3h**) overall inhibited the activity of D1 MSNs, but not D2 MSNs (**Fig. 3i-l**).

### 5HT preferentially inhibits MSNs that signal aggression in the NAc

Next, we took advantage of our cellular-resolution recording technique to determine which D1 MSNs were particularly sensitive to 5HT-mediated inhibition. In two sessions on the same day, we first recorded D1 MSNs during a resident-intruder task, and later recorded the same neurons during optogenetic stimulation of 5HT release. We sorted aggression-modulated D1 MSNs (∼37% of all D1 MSNs) by the magnitude of their GCaMP8m activity at aggression onset (**Fig. 3m**), and then plotted their response to later stimulation of 5HT inputs (**Fig. 3n**). We found that the D1 MSNs that were more active during aggressive behavior were also more inhibited by 5HT release (**Fig. 3o**). In contrast, when we repeated the same experiments with D1-Cre/DAT-Flp mice (**Supplementary Fig. 7a-c**), we found that DA release predominantly excited D1 MSNs, with no preferential effects on aggression-modulated neurons (**Supplementary Fig. 7d-f**). Taken together, our results imply that 5HT, but not DA, selectively inhibits a subset of aggression-active D1 MSNs to suppress attacks. So far we have measured the activity of the same D1 MSNs in two separate sessions: first in a resident-intruder paradigm, and later with optogenetic stimulation of 5HT inputs. However, it is possible that the effect of 5HT on D1 MSN activity differs depending on context. Thus, we ran animals through a third session that combined optogenetic stimulation with the resident-intruder paradigm, allowing us to examine the effect of optogenetically-triggered 5HT release on D1 MSNs during aggression itself (**Supplementary Fig. 8**). Although 5HT release did not change the fraction of D1 MSNs that were activated at aggressive approaches (**Supplementary Fig. 8d**), it significantly suppressed their activity after attack onset (**Supplementary Fig. 8e, f**).

## Discussion

Previous microdialysis measurements of 5HT levels did not have the temporal resolution to capture the sub-second dynamics of 5HT release during aggressive behavior^31^. Here we utilized the latest ‘GRAB’ sensors and optogenetics to reveal that NAc 5HT levels ramp up during aggressive episodes, peaking at aggression offset and helping to terminate attacks. We then combined these manipulations with cellular-level recordings in the NAc, discovering the downstream mechanism of 5HT’s behavioral effect; namely, that 5HT release inhibits a subset of aggression-activated D1 MSNs.

Despite a long history of cross-species work implying that 5HT is anti-aggressive^1,2^, prior studies stimulating all DR neurons or inhibiting all 5HT neurons found little to no acute effect on aggression^32,33^. Our results suggest that DR 5HT neurons may influence aggression in a projection-specific manner, which in this case includes the NAc. Moreover, we found that endogenous NAc 5HT release primarily affected the duration, but not the frequency, of attack bouts. This finding suggests that the NAc may play a specific role in maintaining, rather than initiating, attacks. Closed-loop optogenetics will be an advantageous approach to investigate the temporal specificity and necessity of the DR 5HT-to NAc projection in mediating different phases of aggressive behavior^34–37^.

Using *in vivo* calcium imaging to measure the activity of individual D1 and D2 MSNs, we found that although both populations were activated during aggression, only D1 MSN activity correlated with aggression duration on a trial-by-trial basis. Furthermore, D1, but not D2 MSN inhibition, modulated aggressive behavior. The crucial role for D1 MSNs in aggression was also supported by our experiments in double-transgenic mice, where we stimulated 5HT release and simultaneously recorded the effects of this release on downstream MSN activity. Despite the broad axonal projections and diverse behavioral functions of 5HT neurons^38^, little was known regarding the neuronal response to 5HT release in target areas^8,39^. We found that optogenetically-evoked 5HT release selectively suppressed the activity of D1 MSNs, in particular those that were normally activated during aggression. Furthermore, by recording and stimulating at the same time during the resident-intruder assay, we confirmed that 5HT dramatically suppressed the activity of the aggression-specific D1 neurons. These findings extend previous cFos and chemogenetic results implicating D1 MSNs in aggression reward^21^, adding new evidence supporting the selective engagement of medial shell NAc D1 MSNs in aggressive behavior^21,28,40^. Note, however, that we cannot rule out the possibility that D1 MSN inhibition was triggered by co-release of GABA from 5HT terminals. DR serotonin neurons can co-express gene markers associated with glutamatergic or GABAergic transmission, or both^41,42^. Future work should test whether neurotransmitter or neuropeptide co-release contributes to the neural and behavioral effects observed in our study.

Although we focused much of our work on selective optogenetic stimulation of 5HT inputs, we also performed studies using MDMA, a clinically-relevant, pro-serotonergic compound thought to enhance sociability^18^. We found that systemic injection of MDMA abolished aggressive behavior and preferentially inhibited D1 MSN calcium activity, consistent with both prior behavioral studies^26^ and our own experiments manipulating D1 and D2 MSNs. Previous *ex vivo* slice experiments found that bath application of MDMA reduced excitatory postsynaptic currents (EPSCs) in both D1 and D2 MSNs in NAc slices^18^. The difference between these prior results and the current study are likely due to the myriad differences between *in vitro* and *in vivo* preparations, including the presence versus absence of long-range inputs to the NAc; the 5HT- and DA-releasing potential of MDMA in slices versus intact brains; the precise NAc region that was targeted for recordings; and the differences between recording of evoked EPSCs versus activity-dependent intracellular calcium levels.

Consistent with previous recording^31,43^ and manipulation studies^29,30^, DA release in the NAc increased during aggression but optogenetic stimulation of DA inputs did not modulate aggressive behavior. How did DA release not modulate aggression even though it caused an overall increase in D1 MSN activity? One important difference between the actions of DA and 5HT was that 5HT preferentially inhibited the aggression-active D1 MSNs while DA’s effects did not exhibit such specificity. Thus, the action of DA may leave the relative strength of the aggression-related MSN ensemble unchanged. The mechanisms by which 5HT exerts its inhibitory effect on D1 MSNs remain unknown but likely rely on both presynaptic and postsynaptic 5HT receptors. Presynaptically, we found that infusion of a 5HT1b agonist into the NAc suppressed aggression, presumably in part by suppressing excitatory transmission from glutamatergic inputs to the NAc^17,44^. Postsynaptically, we recently reported that NAc D1 and D2 MSNs express different complements of 5HT receptors^45^ with D1 MSNs expressing higher levels of Gi-coupled (presumably inhibitory) 5HT receptors, while D2 MSNs express more Gq/Gs-coupled receptors. Thus, together, the pre-and postsynaptic actions of 5HT would be expected to inhibit D1 MSNs.

Previous work has shown that 5HT release in the NAc promotes social behavior, as demonstrated by juvenile interaction and three-chamber sociability tests^17,18^. The present findings extend this work by showing that NAc 5HT release also suppresses aggressive behavior through a previously unrecognized mechanism. It is important to note, however, that NAc 5HT release can play multiple other roles. In simple reinforcement learning, for example, NAc 5HT can act in opposition to DA^10^. These observations highlight that the behavioral roles of neuromodulators like 5HT and DA, even within the same structure such as the NAc, are multifaceted and cannot be restricted to single behavioral categories. Nevertheless, by understanding the molecular, cellular, and circuit mechanisms through which 5HT release exerts its pro-social and anti-aggressive effects, we may uncover novel therapeutic strategies for clinical conditions where aggression is a major concern.

## Data and code availability

Data and code are available from the lead contact upon reasonable request.

## Acknowledgements

We thank Daniel F. Cardozo Pinto for useful discussions; Amber Osterman and May Wang for technical support; and Zane C. Norville for advice on the SIMBA software. This work was supported by philanthropic funds donated to the Nancy Pritzker Laboratory at Stanford University and to R.C.M. from the Gatsby Initiative for Brain Development and Psychiatry and the NeuroChoice Initiative of the Wu Tsai Neurosciences Institute. N.E. was supported by NIH grants K08MH123791 and R01MH138645, a Brain & Behavior Research Foundation Young Investigator Grant, a Burroughs Wellcome Fund Career Award for Medical Scientists, a Stanford NeuroChoice Initiative Pilot Award, and a Simons Foundation Bridge to Independence Award. N.E. and R.C.M. are also both supported by ARIA (Aligning Research to Impact Autism). G.C.T. was supported by the Berg Scholars program at Stanford School of Medicine. M.B.P. was supported by NIH grant K99 DA056573. Z.Z. was supported by the Wu Tsai Neurosciences Institute.

## Author contributions

Z.Z, R.C.M., and N.E. conceived the study and designed the experiments. Z.Z. performed the neural recording and optogenetic experiments and analyzed data. Z.Z., G.C.T., P.J.A. and A.N.S. performed the surgeries. T.Y., T.W.H., Z.Z. and P.J.A. performed histology. M.B.P., G.T., and Z.Z. performed the pharmacological experiments. The manuscript was written by Z.Z, R.C.M., and N.E and edited by all authors.

## Methods

### Mice

Wild-type C57BL6/J (Jackson Laboratory #000664), stock number: 017260-UCD), SERT-Flp (Jackson Laboratory #034050), DAT-Flp (Jackson Laboratory #035436), D1-Cre (MMRRC, #029178-UCD) and A2A-Cre (MMRRC, #036158-UCD) mice were used for breeding SERT-Flp^+/-^, D1-Cre^+/-^, A2A-Cre^+/-^, SERT-Flp^+/-^ × D1-Cre^+/-^, SERT-Flp^+/-^ × A2A-Cre^+/-^ and DAT-Flp^+/-^ × D1-Cre^+/-^ mice for neural recording and manipulation. Adult male BALB/c mice (> 8 weeks old, group-housed, 3-5 per cage) were used as intruders into the home cages of male resident mice (> 8 weeks old, singly housed). All experimental mice were housed on a 12-h light/ dark cycle with food and water ad libitum. Behavioral Tests were run in the dark phase. All procedures complied with the animal care standards set forth by the National Institute of Health and were approved by Stanford University’s Administrative Panel on Laboratory Animal Care.

### Surgeries

General surgery procedure: mice (8-12 weeks old, male) were anaesthetized with isoflurane (5% induction, 1.5% maintenance) for surgeries. A stereotaxic frame (Kopf instruments) was used to target injections and implantations (AP and ML are relative to bregma and DV relative to the skull surface). The coordinates used for the NAc were the following in all experiments: AP: +1, ML: 0.7, DV: −4.4 mm. Viruses were infused with a syringe pump (Harvard Apparatus) at a rate of 150 nl/min and allowed to diffuse for at least 5 min. Implants were secured to the skull using screws (Antrin Miniature Specialties) with dental cement (Super-Bond C&B, Sun-Medica and Dual-Cure Resin-ionomer, Geristore). Optical implant placements were histologically verified post-hoc, and mice with mistargeted fibers or lenses were excluded from further analysis. Below lists the virus, target coordinates and implant details specific for different experiments.

For optogenetic stimulation of DR or ventral tegmental area (VTA) terminals, 1μl AAV8-Syn-FLPX-rc [ChRimsonR-tdTomato] (Addgene) or AAV8-Ef1a-fDIO-rsChRmine-oScarlet-WPRE (Stanford viral core) was injected in the DR of SERT-Flp mice, or 0.5 ul of the same virus was injected in bilateral ventral tegmental area of DAT-Flp mice. rsChRmine was used in the optogenetic experiments for its high sensitivity^46^.

For optogenetic inhibition of D1 or D2 MSNs, 700 nl of AAVdj-EF1a-DIO-NpHR3.0-mcherry or AAVdj-Ef1a-DIO-mcherry (Stanford viral core) was injected bilaterally in the NAc of D1-Cre or A2A-Cre mice. Bilateral optical fibers (200-400 µm diameter, Doric) were implanted over the NAc.

For fiber photometry recordings of 5HT or DA, 700 nl of either AAV9-hSyn-5HT3.0 or AAV9-hSyn-GRABDA2m (WZ Biosciences) was injected into the NAc. An optical fiber (400 µm diameter, Doric) was implanted over the NAc for photometry recording.

For ‘Miniscope’ experiments, SERT-Flp × D1-Cre, SERT-Flp × A2A-Cre or DAT-Flp × D1-Cre mice were injected with 1μl of AAV9-syn-FLEX-jGCaMP8m-WPRE (1:2 dilution with phosphate-buffered saline (PBS), Addgene) in the NAc, together with 1μl of AAV8-Syn-FLPX-rc [ChRimsonR-tdTomato] (Addgene) injected in either the DR (AP: −4.6, ML: 0, DV: −3 mm) of SERT-Flp mice or the ventral tegmental area (AP: −3.4, ML: 0.2, DV: −4.25 mm) of DAT-Flp mice. We chose ChRimsonR for the miniscope experiments to minimize potential crosstalk between the blue LED (435-460 nm wavelength for calcium imaging) and optogenetic stimulation^47^. A gradient index (‘GRIN’) lens (0.6 mm in diameter, 7.3 mm long, ProViewTM Integrated Lens, Inscopix) was implanted at the injection site in the NAc.

For drug microinjection, bilateral cannulae (P1 technologies) were implanted into the NAc of mice of wild-type background.

### Fiber photometry

Mice with fiber implants were connected through low-autofluorescence patch cords and a dichroic cube (Doric #FMC6) to a photodetector (Newport #2151). GRAB sensors were excited by frequency-modulated LED light of 405 and 465 nm for near-isosbestic (used as control) and activity-dependent recordings, respectively. The resulting fluorescence signals were demodulated and recorded using a signal processor (RZ5P from Tucker-Davis Technologies running Synapse software) sampled at 1.0173 KHz. For experiments combining fiber photometry with optogenetic stimulation, 9-10 mW, 20 Hz pulses of red LED light (5 ms per pulse) were delivered in either 30-s trains (10 trains, each separated by 90 s; **Fig. 3h, Supplementary Fig. 1k**) or one 3-min train (**Supplementary Fig. 1l**). Raw data was converted into .mat files using software API from Tucker-Davis Technologies (TDT). For the aggression sessions, raw traces were down-sampled by 3x and de-bleached^48^ to get session-wise zscored fluorescence (Fzsc) which was then cropped and aligned to behavioral events (aggression approaches, attacks etc.).

### ‘Miniscope’ imaging and optogenetic stimulation

Calcium imaging and photostimulation were performed using a miniature microscope (nVoke 2.0, Inscopix)^47^. Blue LED light (0.5-1 mW, 435-460 nm) was used for imaging at 10 Hz per imaging plane (512 × 512 pixels per plane). Two planes separated by ∼150 μm were imaged near-simultaneously.

For imaging sessions with MDMA/saline injections, mice were habituated to intraperitoneal injections a day before experiments. On a day of experiment, we recorded baseline activity for 5 min, administered either MDMA (7.5 mg/kg, Sigma-Aldrich) or saline intraperitoneally, waited 5 min, and then recorded for another 20 min. The sequence of MDMA or saline injection was counter-balanced across mice. Injections in the same mice were separated by at least 48 hours.

For imaging sessions during resident-intruder tests and/or optogenetic stimulation, three recordings were completed on the same day with the same field-of-view. The first recording was performed during the resident-intruder test, without any photostimulation. In the second recording, 3 min of photostimulation (12 mW, 20 Hz) was delivered in the middle of a 9-minute resident-intruder assay (with a different intruder). In the third recording, 10 bouts of photostimulation were delivered while the animal was alone in its home cage; these bouts consisted of 30 s, 12 mW, 20 Hz pulses of red LED light (590-650 nm, 5 ms per pulse) separated by 90 s. The order of the first and second recordings was counterbalanced across days, and each mouse experienced two days of experiments (**Supplementary Fig. 8a**).

### ‘Miniscope’ imaging data analysis

Raw imaging data was converted into H5 data files using software from Inscopix (Bruker). Sessions recorded on the same day from the same field-of-view were concatenated before motion correction and neural signal extraction. Piecewise rigid motion correction was applied to each imaging plane^49^. Neuronal shapes, denoised calcium activity traces, and inferred spikes were extracted by a constrained nonnegative matrix factorization (CNMF) method developed for endoscopic datasets^50^. To avoid double counting the neurons that were captured in both imaging planes, the detected ROIs across planes that were spatially close (centroids < 10 pixels apart) and also had high activity correlation (>0.5) were merged into one. The calcium traces were z-scored for each session and aligned to either behavior timestamps (aggressive approach, attack etc.) or photostimulation onsets. These z-scores were then baseline-corrected to produce ΔFzsc, with the baseline defined as either the 1 s window starting 2 s before the behavior, or the 10 s window before photostimulation onset. Timestamps within 3 s of a prior behavioral event were ignored to avoid confounding baseline subtraction with behavior-related activity.

To identify aggression-modulated neurons, the aggression-aligned activity (averaged in a 2 s window following each aggressive approach) was compared to a distribution of activity values aligned to 1000 random timestamps. A neuron was considered to be aggression-modulated if its aggression-aligned activity was higher or lower than 90% of the random-timestamp-aligned activity values. For sessions with drug or saline injections, the inferred spike rate from each neuron was normalized by that during the baseline period.

### Optogenetic manipulations during resident-intruder

For optogenetic manipulations, mice with optical implants were connected to a 625 nm LED light source (Prizmatix) via a plastic fiber (1 mm diameter, NA 0.63) and a fiber optic rotary joint (Doric). For photostimulation of DA or 5HT inputs to the NAc, mice underwent two days of optogenetics experiments. On each day, they performed two 9-minute sessions separated by >1h, one with photostimulation and one without. The order of photostimulation was counterbalanced between days. The photostimulation (3 min at 20 Hz, 12 mW, 5 ms pulse width) was initiated 3 min after intruder entry. For photoinhibition of D1 or D2 MSNs in the NAc, mice likewise underwent two sessions, separated by >1 hour. In the second session, we delivered a 9-minute train of 0.4 Hz, 8 mW, 2 s light pulses. The duration of optogenetic terminal stimulation was chosen to balance two factors: it had to be long enough to capture enough aggressive bouts to test the behavioral effect, but short enough to avoid opsin desensitization. Similar durations of 5HT terminal stimulation (5-min for three-chamber sociability and 2-min for juvenile interaction) produced pro-social effects^17^, so we hypothesized that 3-minute elevations of 5HT levels in the NAc would similarly influence attacking behavior. Video recordings and optogenetic stimulations were synchronized by TTL triggers from a data acquisition card (USB6259, National Instrument) controlled by PackIO^51^.

### Drug injections

For intracranial drug microinjection, two 9-minute sessions were performed, separated by >1 h. 15 min before the second session. On one test day, after the first session. mice received a bilateral intra-NAc infusion of either saline with 1% DMSO, or one of the selective 5HT receptor agonists: 0.5 µg CP93129 diluted in saline targeting 5HT1b receptors; 1 µg LY344864 diluted in 1% DMSO targeting 5HT1f receptors, or 0.5 µg WAY163909 diluted in saline targeting on 5HT2c receptors. Drugs were infused through an injector cannula using a microinfusion pump (Harvard Apparatus) at a continuous rate of 0.1 µl per min to a total volume of 0.3 µl per hemisphere. Injector cannulae were removed 2 min after infusions were complete. Injections in the same mice were separated by at least 48 hours.

For systemic MDMA/saline injections, mice were habituated to intraperitoneal injections a day before experiments. On one day. either MDMA (7.5 mg/kg, Sigma-Aldrich) or saline was administered intraperitoneally 15 min before the behavioral tests. The sequence of MDMA or saline injection was counter-balanced across mice. Injections in the same mice were separated by at least 48 hours.

### Resident-intruder tests

Mice were habituated to the recording arena (with fiber connection for optogenetic experiments) by performing a daily resident-intruder test until they attacked the intruder within 5 min after the intruder was introduced, and more than 2 times in 9 min on two consecutive days. Mice for microinjection experiments underwent a sham injection handling before these habituation sessions. 32 mice that did not reach the aggression criteria after 1 week were excluded from further experiments. On rare occasions when an intruder was aggressive towards the resident, the experiment was immediately aborted, and the intruder was not used for further experiments. For each test mouse, a different intruder was used for control (saline or no-light) and test (drug-treated or light-on) conditions. All resident-intruder sessions were 9-min long. We did not observe any injuries to the intruder that would require capping the number of bites or ending the session early. For detailed descriptions of how we combined resident-intruder tests with imaging or manipulations, please see the relevant sections above.

### Resident-intruder behavior analysis

Behavioral chambers were illuminated with dim red light and videos were taken by an overhead camera (FLIR Blackfly) at 20 Hz frame rate in SpinView software (Teledyne Vision Solutions). Mouse postures were extracted from behavior video recordings using ‘DeepLabCut’^52^. For sessions with neural activity recordings, to get the exact onset time of attacks, videos were manually labelled by experimenters who were blinded to experimental condition or simultaneously-collected imaging data. To obtain more objective timestamps of behaviorally-relevant epochs in our neural data, we defined the onsets/offsets of aggression episodes (i.e. the ‘aggressive approaches’ and ‘aggression offsets’) as the time points when the two mice became close to (< 5 cm) or far away from (> 5 cm) each other before/after attacks. To reduce confounds in our baselining, we only analyzed bouts that lasted at least 2 s and were separated by at least 2 s from a prior bout. For sessions without neural activity recordings, we used a machine learning based software, ‘SIMBA’^53^, to identify attacking behavior (**Supplementary Notes, Supplementary Video 1**). The detected attacking bouts that were < 0.25 s apart were merged into one, and those that were shorter than 0.5 s were filtered out. The behavioral metrics (duration per attack, sniffs per minute etc.) under LightOff and LightOn conditions were divided by the corresponding value in the LightOff condition to get a percentage change (%).

### Open field test

Mice were placed into a 40 x 40 cm behavioral arena and allowed to explore freely for 12 mins. Optogenetic manipulations were delivered in four alternating light-on (0.4 Hz, 8 mW, 2 s light pulses for NpHR3.0 stimulation) and light-off epochs that lasted 3 mins each. Epoch order was counterbalanced across mice. Biobserve Viewer behavioral tracking program was used to extract distance traveled during each epoch.

### Real-time place preference

Real-time place preference (RTPP) tests were performed in an arena with three chambers as previously described^10^: a neutral center chamber with clear floors, and two identical side chambers with white walls and floors. Mice were confined to the center chamber for a ∼2 min habituation period. Then, the barriers were removed to begin the first half of the test during which mice were free to explore all three chambers for 15 min and one side chamber (left or right, counterbalanced) was paired with photostimulation (20 Hz, 12 mW, 5 ms pulses for terminal ChRimsonR or rsChRmine stimulation, or 0.4 Hz, 8 mW, 2 s light pulses for cell-body NpHR3.0 stimulation) that began when the mouse entered the light-paired chamber and continued for the entire duration that the mouse occupied the chamber. After 15 min, the side that was paired with photostimulation was swapped and mice were free to explore all three chambers for an additional 15 min reversal phase.

### Three-chamber sociability assay

A three-chamber sociability assay was performed in an arena with three separate chambers as previously described^17^. On day one, the test mice were habituated to the arena, with two empty wire mesh cups placed in the two outer chambers for 5 min. On day two, the test mouse was placed in the center chamber and a conspecific juvenile (3–5-week-old males) was placed under one of the wire mesh cups. The test mice were first placed in the center chamber for 2 min. The barriers were then raised and the test mouse was allowed to explore freely for a 20-min session with 5-min epochs of photostimulation (0.4 Hz, 8 mW, 2 s light pulses for cell-body NpHR3.0 stimulation). The order of these epochs (LightOn or LightOff) was counterbalanced across mice. The placement of juvenile mice in the chamber was also counterbalanced across sessions.

### Histology

Mice were transcardially perfused with 4% (w/v) paraformaldehyde (PFA) in phosphate-buffered saline (PBS) and postfixed in PFA overnight. Coronal sections 70 µm thick were immunostained for GFP (primary: chicken anti-GFP, Aves 608 #GFP-1020; secondary: goat anti-chicken 488, Invitrogen #A-11039), and/or mCherry (primary: rat anti-mCherry, Invitrogen #M11217; secondary: goat anti-rat 594, Invitrogen #11007). Primary antibodies were used at a concentration of 1:1000 and incubated on a shaker overnight at room temperature. Secondary antibodies were used at a concentration of 1:750 and incubated on a shaker for 2 hours at room temperature.

### Statistics and reproducibility

Statistical analyses were performed in MATLAB (MathWorks) and Python. No statistical methods were used to predetermine sample sizes, which were based on previous experience with the variance of the assays. Statistical parameters such as the types of statistical tests, sample numbers (n) and statistical significance are reported in the figures and figure legends. T-tests were used to test if a sample mean was different from zero. Wilcoxon rank-sum tests were performed to compare unpaired measurements between two groups, and Wilcoxon signed-rank tests were conducted to compare paired measurements. For multi-group comparisons, we conducted one-or two-way analyses of variance followed by Tukey’s honestly significant difference procedure. All statistical tests were two-sided. All error bars in the figures are s.e.m. unless otherwise noted.

---

**Supplementary Figure 1:**
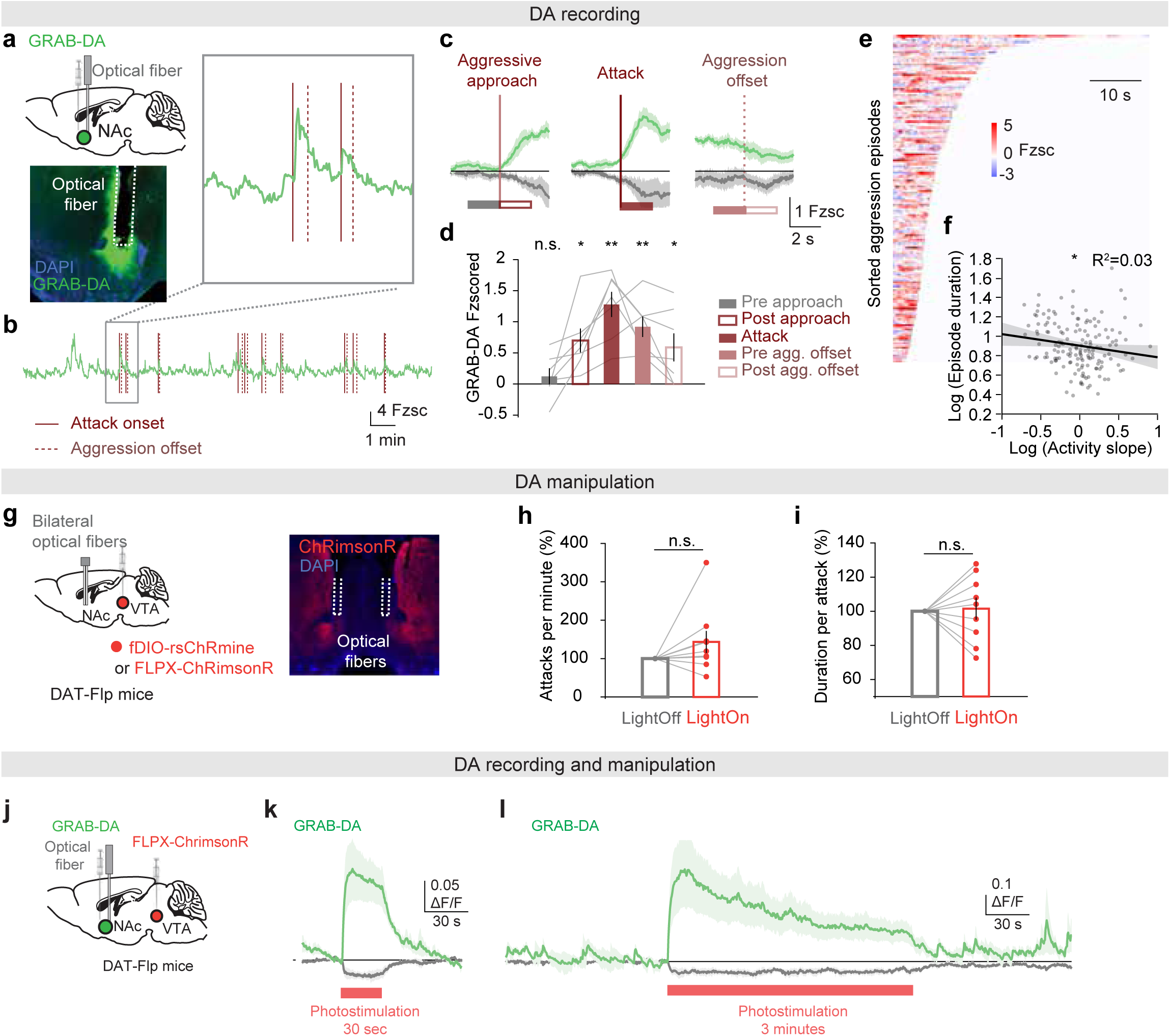
DA signaling in the NAc during aggression. (a) GRAB-DA virus was injected in the NAc for fiber photometry recording. (b) Example session of GRAB-DA recording during a resident-intruder test. (c) Average GRAB-DA (green) and control (grey) signal aligned to aggressive approach, attack, or aggression offset. (d) GRAB-DA signal averaged in a 2-s window pre/post the behavioral events as marked in (c) (left to right: n.s. P = 0.38, *P= 0.011, **P = 0.001, 0.001, *P = 0.040, one-sample t-test, n = 7 mice). (e) Heatmap showing GRAB-DA signaling in individual aggressive bouts sorted by duration. (f) The duration of aggressive episodes negatively correlates with the slope of DA increase (*P=0.042, r = −0.16, Pearson’s correlation coefficient). (g) Flp-dependent excitatory opsin was injected into the ventral tegmental area of DAT-Flp mice. A bilateral optical fiber was implanted in the NAc for optogenetic stimulation of DA terminals. (h) Change in the frequency of attacks in photostimulation compared with non-photostimulation sessions (n.s. P = 0.16, Wilcoxon signed-rank test, n = 9 DAT-Flp mice). (i) Same as (h) but for the change in average duration of attack bouts (n.s. P = 0.82, Wilcoxon signed-rank test). (j) Flp-dependent ChRimsonR virus was injected in the VTA, while GRAB-DA virus was injected in the NAc of DAT-Flp mice. (k-l) GRAB-DA signal during photostimulation of ChRimsonR-expressing VTA DA terminals (n = 3 mice, shaded areas show s.d.).

**Supplementary Figure 2:**
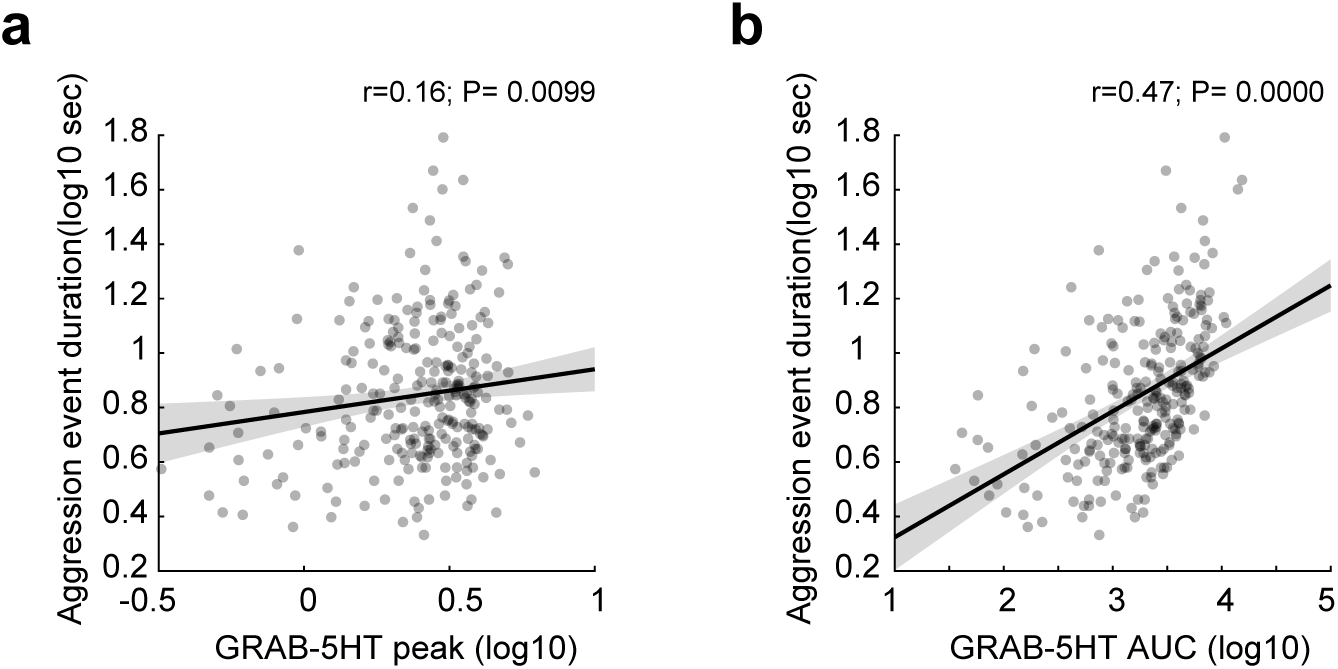
Other metrics of 5HT release vs aggression duration. Peak (a) and area under curve (AUC, b) of GRAB-5HT signal during each aggression episode plotted against the episode durations (same dataset as in Figure 1, Pearson’s correlation coefficient).

**Supplementary Figure 3:**
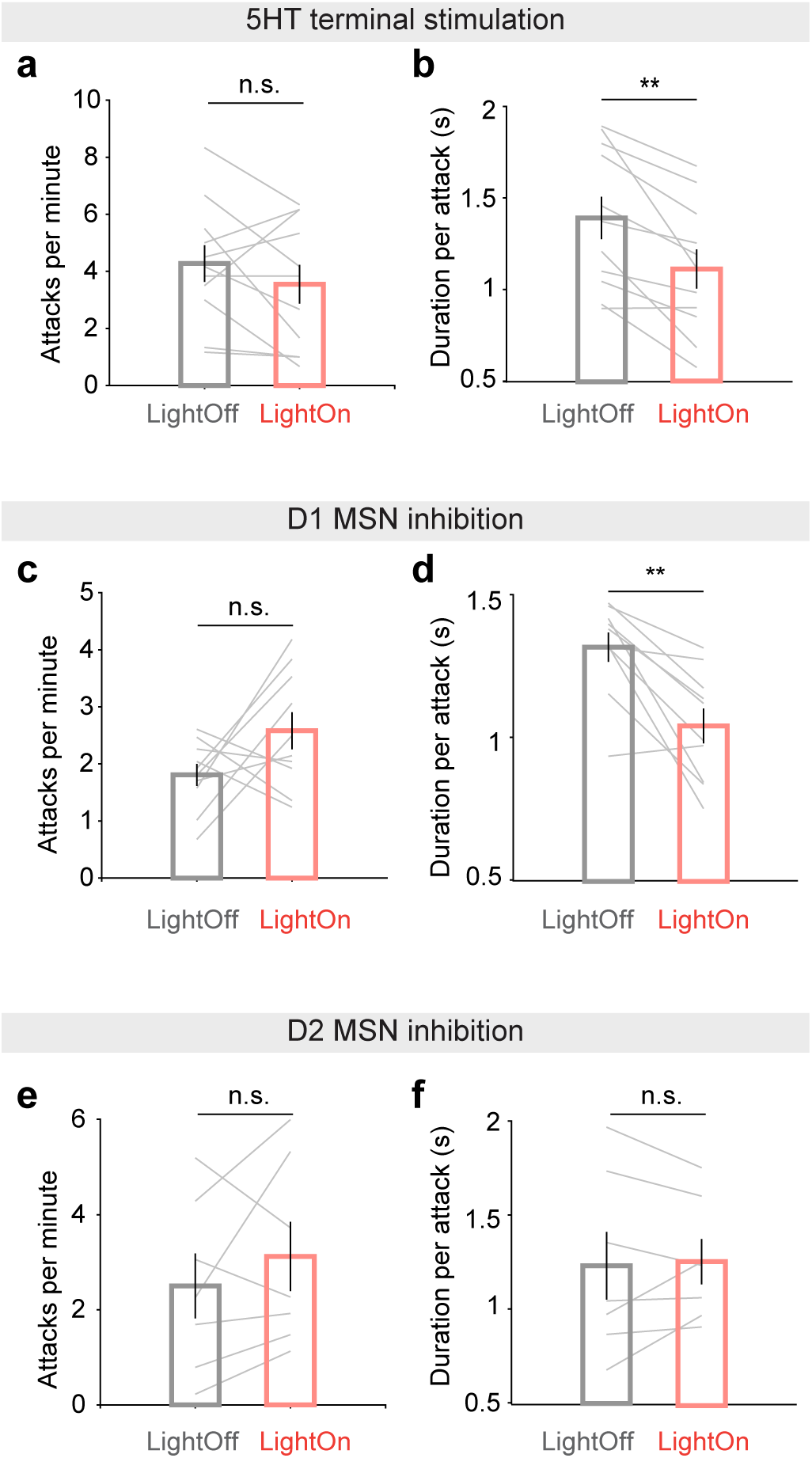
Raw behavior measurements during resident-intruder tests. Frequency and duration of attacks in LightOn and LightOff conditions during 5HT terminal stimulation (a-b, n.s. P = 0.28, ** P = 0.0020), D1 MSN inhibition (c-d, n.s. P = 0.16, ** P = 0.0039) and D2 MSN inhibition (e-f, n.s. P = 0.38, 0.81, Wilcoxon signed-rank test). Same dataset as in Figure 1j,k and Figure 2h,i, displaying raw instead of normalized values.

**Supplementary Figure 4:**
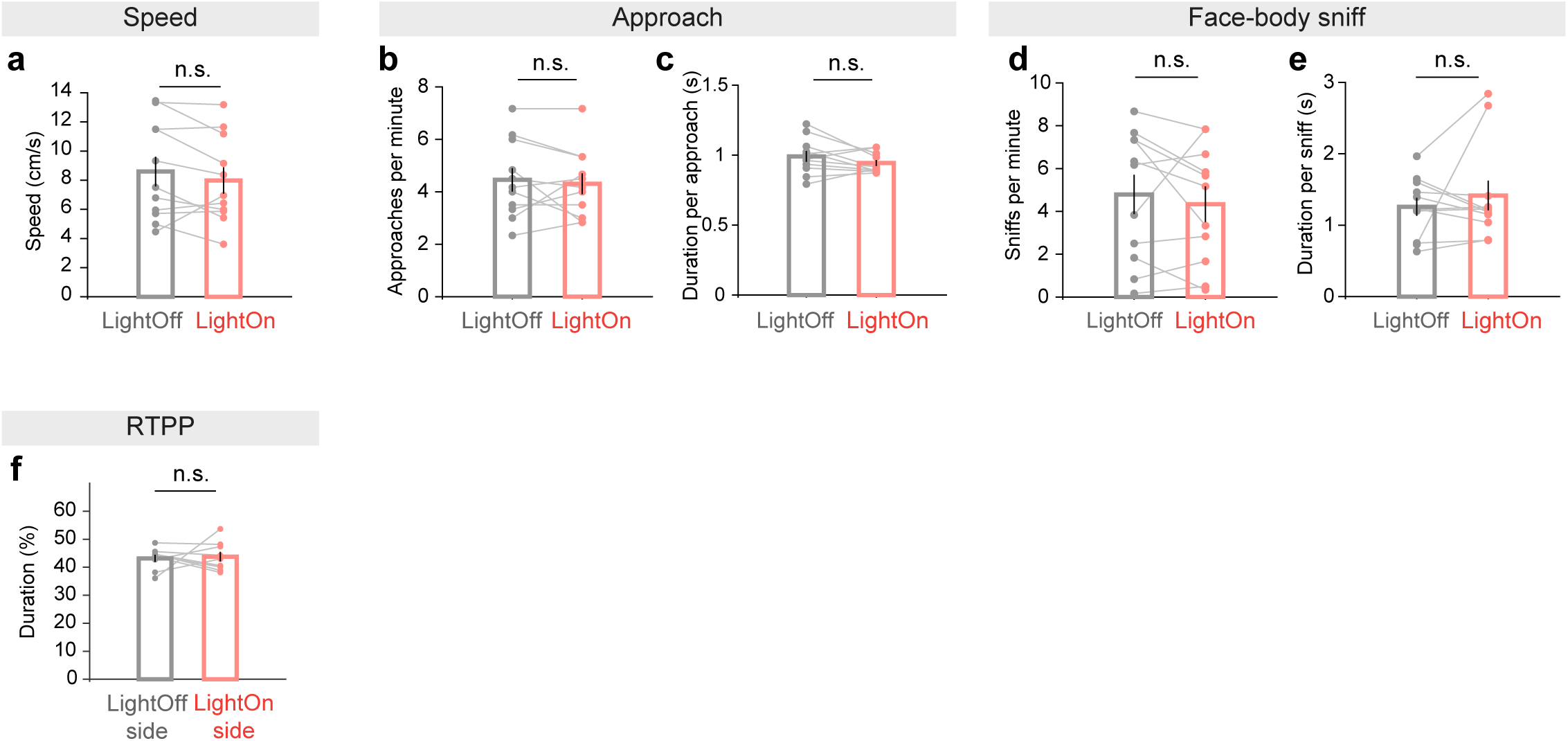
Non-aggressive behaviors during SERT terminal stimulation. Change in running speed (a) frequency and duration of social approaches (b-c) or face-body sniffs (d-e) with and without 5HT terminal stimulation (n.s. P>0.05, Wilcoxon signed-rank test, same mice as in Figure 1j-k) (f) Fraction of time spent in the stimulation-paired chamber (‘LightOn side’) and the non-paired chamber (‘LightOff side’) in the real-time place preference (RTPP) tests.

**Supplementary Figure 5:**
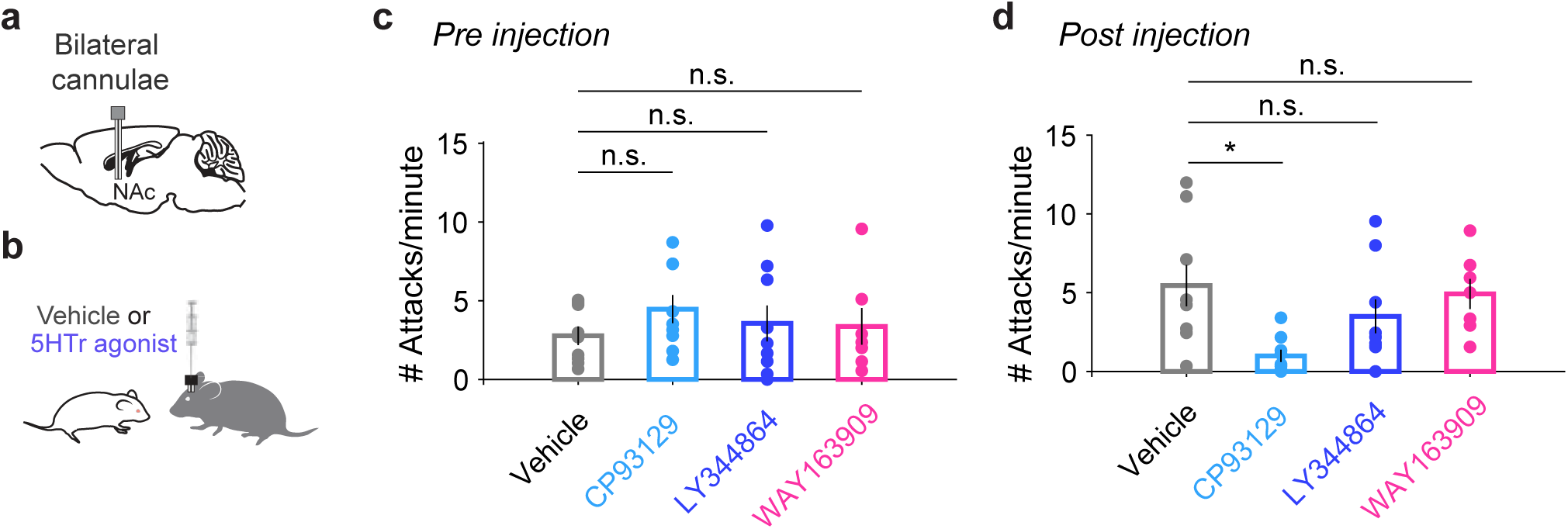
Local infusion of 5HT1b receptor agonist in the NAc significantly suppressed aggression. (a) Bilateral cannulae were implanted over the NAc for microinjection of either vehicle solution alone or with a selective 5HT agonist. (b) One type of 5HT receptor agonists or vehicle solution was injected in the NAc before resident-intruder tests. (c) Frequency of attacks in a baseline session 1 hour before drug injection. (d) Frequency of attacks 15 minutes post microinjection of either vehicle alone (1% DSMO, n = 8 mice), 5HT1b receptor agonist (CP93129, n = 8 mice), 5HT1f receptor agonist (LY344864, n = 7 mice) or 5HT2c receptor agonist (WAY163909, n = 6 mice). n.s. P>0.05, *P < 0.05, one-way ANOVA followed by Tukey-Kramer test.

**Supplementary Figure 6:**
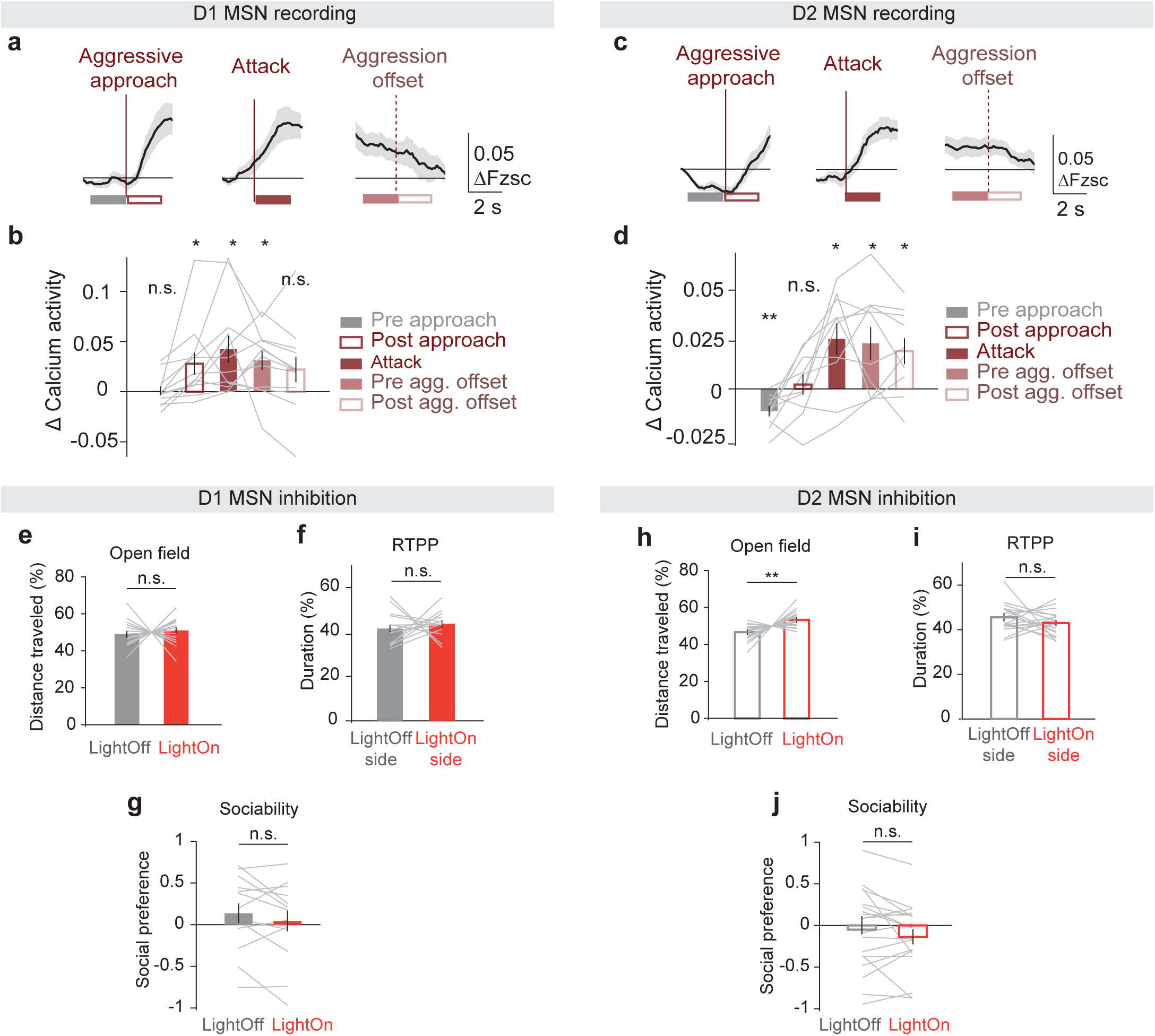
MSN recording and manipulation in the NAc. (a) Average GCaMP signal of D1 MSNs aligned with aggressive approach, attack, or aggression offset. (b) Calcium activity averaged in a 2-s window pre/post the behavioral events as marked in (a) (left to right: n.s. P = 0.86, *P = 0.027, 0.010, 0.006, n.s. P = 0.12, one-sample t-test, n = 12 mice). (c-d) Same as (a-b) but for D2 MSNs (left to right: **P = 0.002, n.s. P = 0.067, *P = 0.011, 0.021, 0.017, n = 10 mice). (e) Distance traveled during the light-on and light-off epochs in open field tests (n.s. P = 0.96, n = 16 D1-Cre mice injected with DIO-NpHR3.0 in bilateral NAc and implanted with optical fibers over injection sites). (f) Duration of time spent on each side of the chamber during RTPP tests (n.s. P = 0.35, n = 14 D1-Cre mice). (g) Social preference score during LightOff and LightOn periods in three-chamber sociability tests (n.s. P = 0.27, n = 14 D1-Cre mice). (h-j) same as (e-g) but for D2 MSN inhibition (**P = 0.0038 in h, n.s. P = 0.30 in i, n.s. P = 0.24 in j, n = 19 A2A-Cre mice, Wilcoxon signed-rank test).

**Supplementary Figure 7:**
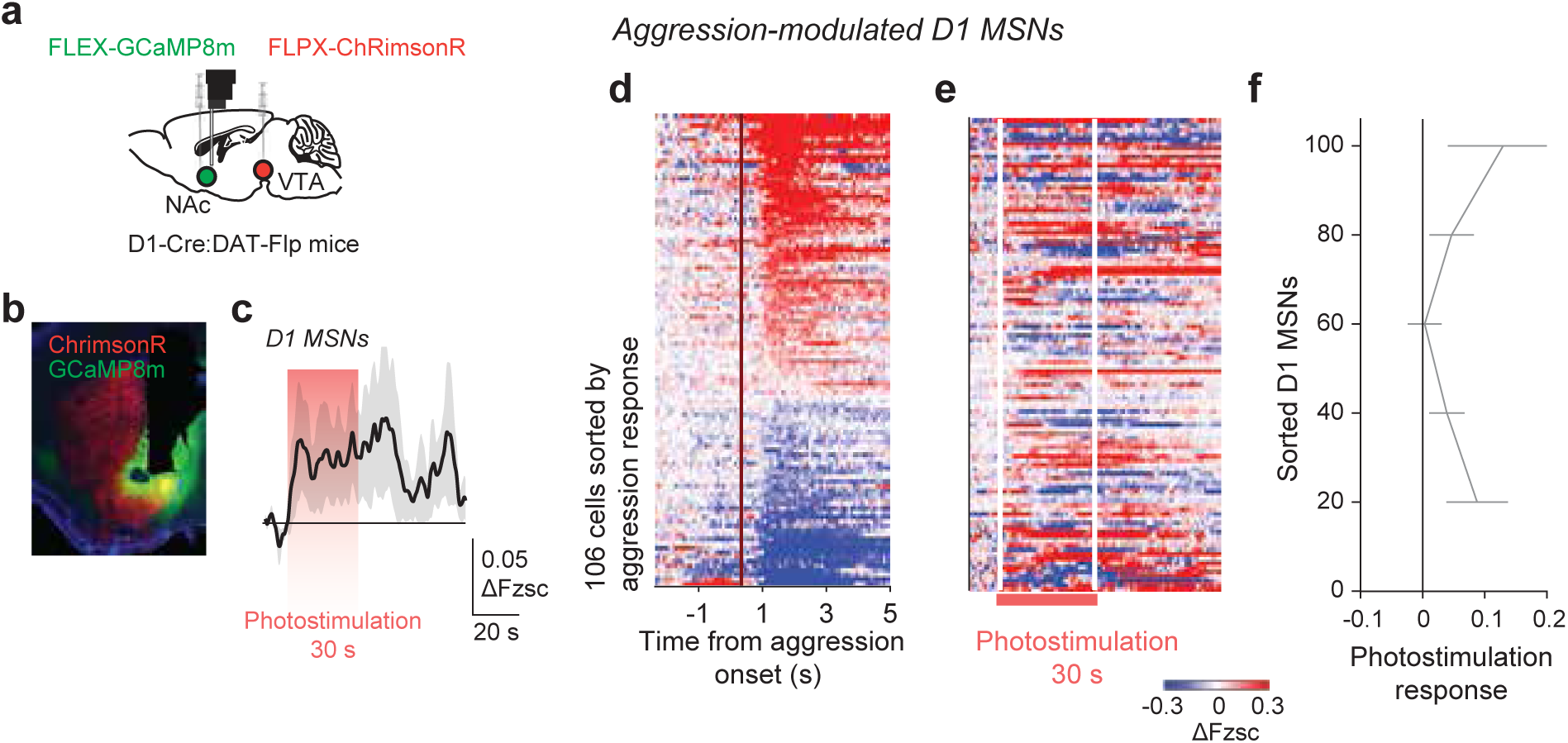
Optogenetic activation of VTA DA terminals while imaging D1 MSNs in the NAc. (a) Experimental strategy: Flp-dependent ChrimsonR virus was injected in the VTA, while Cre-dependent GCaMP8m virus was injected in the NAc of D1-Cre DAT-Flp mice. (b) Example brain slice of the NAc with GCaMP8m expressed in D1 MSNs and ChrimsonR in DA terminals. (c) Calcium activity of D1 MSNs aligned to photostimulation (30 s, 30 Hz) of VTA DA terminals. (d) Calcium activity of aggression-modulated D1 MSNs around aggressive approach. Each row is a cell, sorted by the calcium response averaged between 0-2 s after aggression onset (n = 8 sessions, 4 mice). (e) Calcium activity of the same cells in (d) aligned with photostimulation. (f) Photostimulation response of cells sorted by aggression response as in (d-e).

**Supplementary Figure 8:**
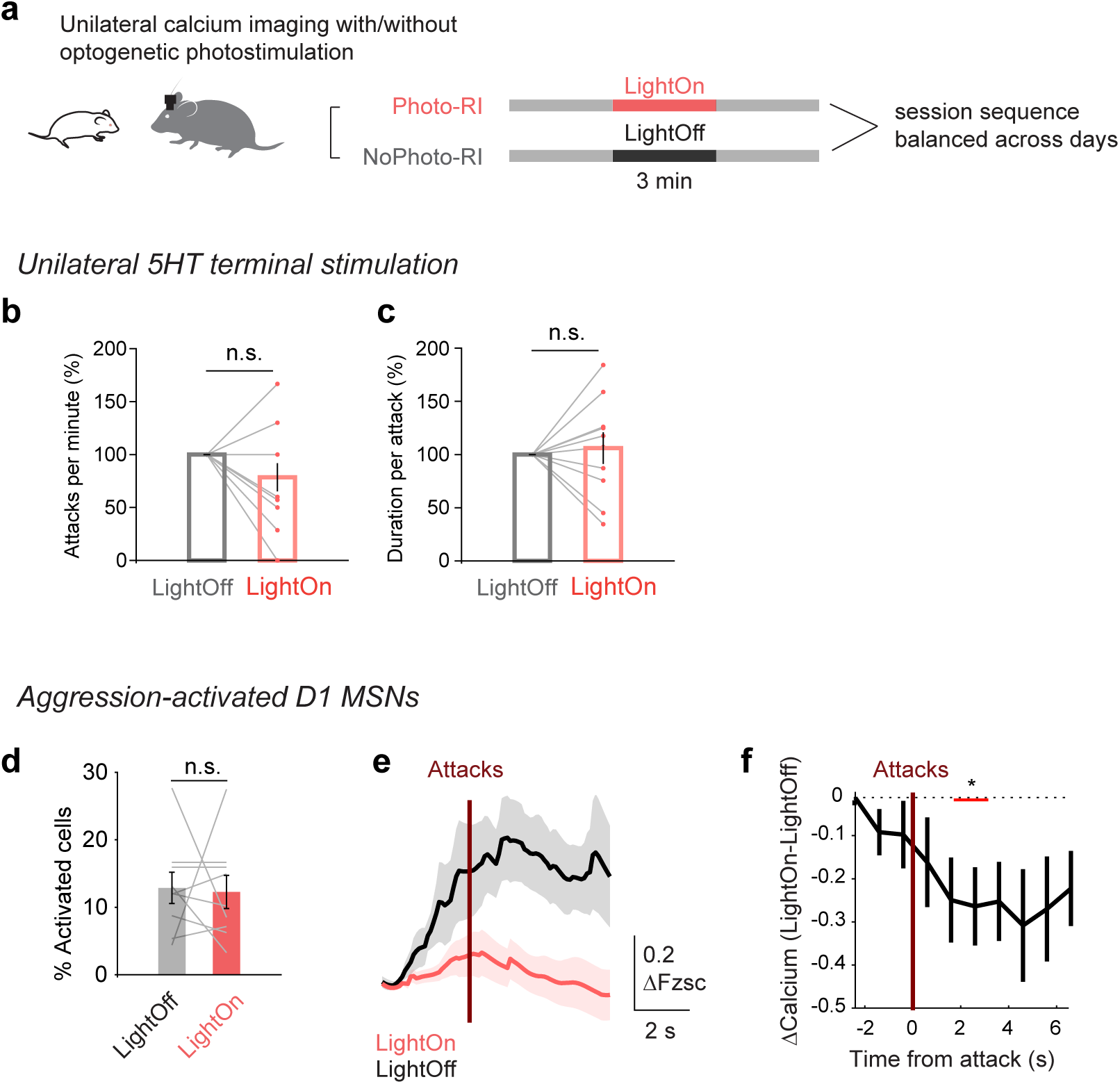
Aggression-activated D1 MSNs are inhibited by 5HT terminal stimulation during the resident-intruder test. (a) Mice performed the resident-intruder (RI) assay twice in a single day: once with photostimulation (Photo-RI) and once without (NoPhoto-RI). The sequence of Photo-RI and NoPhoto-RI sessions was balanced across days. (b-c) Frequency and duration of attacks in LightOn and LightOff conditions during miniscope experiments where 5HT terminals were stimulated unilaterally (n.s. P = 0.13, 0.62, Wilcoxon signed-rank test, n = 6 D1-Cre × SERT-Flp mice and n = 6 A2A-Cre × SERT-Flp mice). (d) Numbers of D1 MSNs that were activated at aggressive approaches during LightOn and LightOff episodes (n.s. P = 0.81). (e) Average activity of aggression-activated D1 MSNs aligned to attacks. (f) Difference in the LightOn and LightOff activity traces in (e) binned in 1-s bins (* P = 0.047, Wilcoxon signed-rank test, n = 7 sessions with attacks in both LightOff and LightOn episodes).

## DLC-SIMBA pipeline for resident-intruder video analysis

We used the classic DeepLabCut (DLC) method^1^ without multi-animal tracking functionality to extract poses of the test mice and the intruders, which are of distinct colors (dark gray vs white, **Supplementary Video 1**). The training dataset details are summarized in **Table 1**. Eight body parts (‘keypoints’) were used for each animal: Ear_left, Ear_right, Nose, Center, Lateral_left, Leteral_right, Tail_base, and Tail end. The model achieved train error of 2.13 pixels and test error of 2.18 pixels after 675,000 iterations. We found that the ‘Tail end’ point was often off in test videos, which presumably contributed most to the errors, and so these keypoints were not used for further behavioral identifications.

**Table 1:** Training dataset characteristics for generating a DeepLabCut model.

| Feature | Value |
| --- | --- |
| Keypoints (total) | 16 |
| Individuals | 2 |
| Labeled frames | 522 |
| Labeled videos | 20 |
| Total duration (s) | >10800 |

We used three previously-trained SIMBA classifiers^2^ for detecting attacks, approaches, and sniffs (face to body). Approaching and face-body sniffing were used as less-aggressive control behaviors. The training sets are summarized in Table 2. The behavioral definitions are summarized in Table 3.

**Table 2:** Training sets for the three SIMBA classifiers.

| Classifier | No. annotated frames | No. of positive frames | No. of videos | Optimal threshold | F1 | Precision | Recall |
| --- | --- | --- | --- | --- | --- | --- | --- |
| Attack | 126604 | 25214 | 27 | 0.7665 | 0.868 | 0.798 | 0.951 |
| Approach | 126604 | 5881 | 27 | 0.7689 | 0.836 | 0.747 | 0.948 |
| FaceBody sniff | 126604 | 26958 | 27 | 0.8269 | 0.839 | 0.771 | 0.912 |

**Table 3:** Operational behavioral definitions for SIMBA classifiers.

| Classifier | Description | Start frame | Duration of behavior | End frame |
| --- | --- | --- | --- | --- |
| Attack | Clear physical antagonistic interaction initiated by the Resident mouse. Brief pauses can occur if the Resident mouse remains oriented toward the Intruder. | Resident mouse makes physical antagonistic contact with the Intruder. Typically this is characterized by outstretched Resident forepaw(s) contacting the Intruder, while the Resident has an open mouth to initiate a bite. Can also be characterized by the first frame of a slap or quick bite without the forepaws being outstretched. | Short breaks may be present in attack behavior, if resident is still oriented toward Intruder. Attacks can include tussling, biting, boxing, and corralling as part of the attack bout. | Resident mouse orients away from Intruder. Typically this is a slight turning of the head to look in a different direction, followed by a relaxation of the body and moving away from the Intruder. |
| Approach | Resident mouse orients self to point at Intruder and moves directly toward Intruder. Intruder is not actively moving away from Resident and may be either stationary or engaging another non-escape behavior, including grooming and operant chamber exploration. | Resident mouse is moving directly towards a non-escaping intruder. | Resident is oriented at and moving directly toward Intruder. Intruder is not moving directly away from resident and may be engaging in non-escape behaviors. The Intruder may still traverse the operant chamber so long as it does not begin specifically moving away from an approaching resident. | Resident mouse stops moving toward anon-escaping intruder mouse or when intruder begins moving directly away from resident. |
| FaceBody sniff | Resident mouse is sniffing the Intruder, but not in the anogenital region (base of tail). | Resident mouse is clearly sniffing the intruder in a facial or bodily region | Uninterrupted sniffing of facial or bodily region | Resident mouse moves head away from Intruder, sniffs the anogenital/tail region, or engages in a separate behavior. |

To re-use the classifiers for new videos captured by a different camera lens, we scaled our DLC outputs by changing the SIMBA input parameter, ‘pixels/mm’, from 1.51 (used for the training set) to 0.9. We used threshold = 0.8 for all three behaviors. To avoid continuous behavioral bouts being incorrectly separated into multiple bouts, or false positive detections in single frames, the detected bouts that were < 0.25 s apart were merged into one, and those that were shorter than 0.5 s were filtered out.

Finally, to validate the performance of the full DLC-SIMBA pipeline, we annotated five videos recorded under our experimental conditions, which were not used for training either the DLC or the SIMBA models. Using the manual labels as ‘ground truth’, the performance of the full pipeline is summarized in Table 4.

**Table 4:** Performance of classifiers on test videos.

| Classifier | F1 | Precision | Recall |
| --- | --- | --- | --- |
| Attack | 0.712 | 0.617 | 0.842 |
| Approach | 0.647 | 0.719 | 0.517 |
| FaceBody sniff | 0.602 | 0.640 | 0.654 |

Plotting hand annotation against DLC-SIMBA detection results (**Figure R1**), we found that although there are offsets between the values, as reflected in the shifts or tilts of the linear fits (red) from the diagonal lines (grey), the hand annotation and SIMBA detection results, in particular for attack and approach, are significantly and linearly correlated, with R’s>0.8. The offsets may in part reflect the annotator’s bias (as she was not involved in annotating the training data).

**Figure R1:**
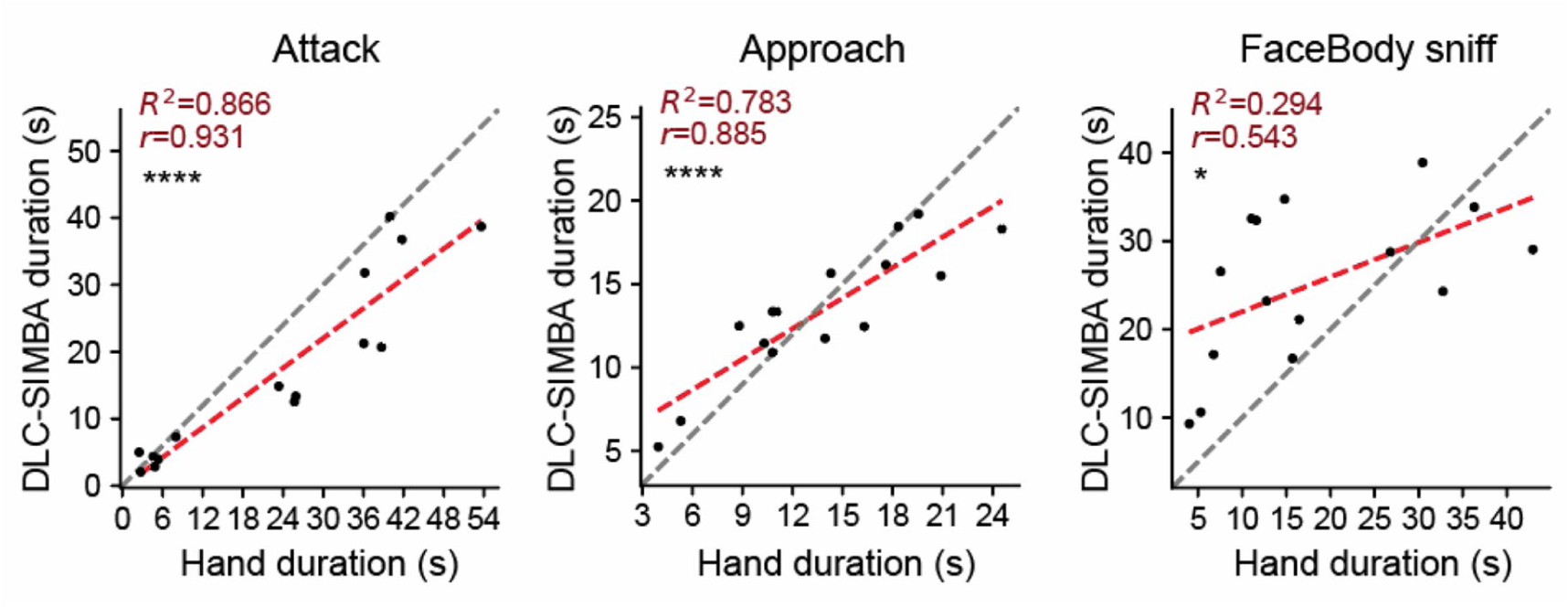
Hand annotation versus DLC-SIMBA detections for each behavior. (****P<0.0001, *P = 0.036, n = 5 videos, 5 resident-intruder pairs; each 9-min video was cut into 3 x 3-min sections and each dot corresponds to the duration of a specific behavior in a section)

We used the machine learning pipeline to analyze videos in optogenetic/drug manipulation experiments, where the relevant outcome was a behavioral change between test and control conditions. We preferred the machine learning pipeline in these analyses to avoid human bias in comparing behavioral results across conditions. In contrast, for the neural recording experiments, we wanted to maximize timing accuracy so that we could align the neural activity to behavioral timestamps on a frame-by-frame basis. We avoided bias by blinding the experimenter from the neural data while they labeled frames. We felt that manual labeling, although time-consuming, would maximize the accuracy of timestamps for neural data analyses.

